# Characterization of HcaA, a novel autotransporter protein in *Helicobacter cinaedi*, and its role in host cell adhesion

**DOI:** 10.1101/2022.11.18.517170

**Authors:** Sae Aoki, Shigetarou Mori, Hidenori Matsui, Keigo Shibayama, Tsuyoshi Kenri, Emiko Rimbara

## Abstract

*Helicobacter cinaedi* infects the human gut and causes invasive infections such as bacteremia and cellulitis through bacterial translocation. However, the mechanism by which *H. cinaedi* attaches to host cells and establishes infection remains unclear. This study aimed to investigate the relationship between a novel putative autotransporter protein, *H. cinaedi* autotransporter protein A (HcaA), and its role in pathogenicity. The cytotoxicity of *H. cinaedi* infection in colon epithelial cell lines (Caco-2 and HT-29) was assessed using a lactate dehydrogenase assay, and it was found that cytotoxicity significantly decreased upon HcaA knockout. Adhesion assays further revealed that the HcaA-knockout strain showed significantly reduced attachment to the human epithelial colorectal adenocarcinoma cell line (Caco-2) compared to that of the wild-type strain. Moreover, the recombinant HcaA protein demonstrated strong adhesion properties to the human monocytic cell line (U937). The adhesive activity was diminished when the RGD motif in HcaA was replaced with RAD, indicating that the RGD motif in HcaA is crucial for host cell adhesion. To determine the role of HcaA in *H. cinaedi* infection *in vivo*, C57BL/6 mice were orally infected with wild-type and HcaA-knockout *H. cinaedi* strains. Bacterial colonization was assessed 7, 14, and 28 days post-infection. At 7 days post-infection, colonization was significantly lower in mice infected with the HcaA-knockout strain compared to those infected with the wild-type strain. In conclusion, our findings suggest that HcaA, a novel putative autotransporter protein in *H. cinaedi*, plays a significant role as an adhesin in establishing colonization.

**IMPORTANCE:** *Helicobacter* species are classified as gastric or enterohepatic according to their habitat. Among enterohepatic *Helicobacter* species, which inhabit the intestine, colon and liver, *H. cinaedi* has been most frequently isolated from humans. *H. cinaedi* often causes bacteremia and cellulitis in immunocompromised hosts. Here, we focused on the *H. cinaedi* autotransporter protein A (HcaA), a novel virulence factor in *H. cinaedi*. We discovered that HcaA contributes to cell adhesion via its RGD motif. Furthermore, in animal experiments, bacterial colonization was reduced in mice infected with HcaA-knockout strains, supporting the hypothesis that HcaA contributes to *H. cinaedi* adhesion to host cells. Our study provides a novel mechanism for the establishment of *H. cinaedi* infections and provides new insights into the role of autotransporter proteins in the establishment of *Helicobacter* infection.

## INTRODUCTION

*Helicobacter* species are gram-negative, helical bacilli isolated from humans and mammals. They are classified into gastric or enterohepatic *Helicobacter* species based on their habitat. *Helicobacter pylori*, a typical gastric *Helicobacter* species, infects the human stomach and is a major cause of gastric cancer (1). *H. cinaedi* is the most frequently isolated enterohepatic *Helicobacter* species strain from humans, mainly found in immunocompromised patients with bacteremia. *H. cinaedi* is responsible for recurrent cellulitis, infected aortic aneurysm, subdural hematoma, and hepatic cyst infection (2–6). *H. cinaedi* was first isolated in 1984 from a homosexual man with intestinal symptoms in Seattle and it was first isolated in Japan in 2003 (7, 8). Though *H. cinaedi* infections are predominantly reported in Japan, there have been some case reports from Denmark (9, 10). A retrospective study at a single hospital in Japan indicated that, 2.2 % of all blood culture-positive samples turned out to be *H. cinaedi*-positive when the blood culture period was extended, suggesting that *H. cinaedi* infections might be overlooked in other countries (11).

Cytolethal distending toxin (CDT) is a virulence factor in a variety of gram-negative pathogenic bacteria, including *H. cinaedi*. It consists of three polypeptide subunits, CdtA, CdtB, and CdtC (12–14). CdtB possesses nuclease activity and causes DNA damage, leading to senescence and apoptosis in host cells. CdtA and CdtC play a role in delivering CdtB into host cells. Additionally, another virulence factor, *H. cinaedi* antigen CAIP (15), causes the differentiation and maintenance of the pro-inflammatory profile of human macrophages and triggers foam cell formation. Consequently, inflammation in atherosclerosis is induced. Studies have been conducted on *H. cinaedi* biology and host-host interactions protection against oxidative stress and changes in virulence via the production of alkyl hydroperoxide reductase (16) as well as changes in oxygen demand during infection (17). However, the mechanisms by which *H. cinaedi* interacts with host cells and establishes infection have not been fully elucidated.

Autotransporter proteins (ATs) are type V secretion systems (T5SSs) comprising an essential passenger domain that functions as the secreted portion, and an autotransporter domain with a β-barrel that exports the passenger domain from the periplasm to the extracellular environment. ATs are widely found in gram-negative bacteria and have diverse functions, acting as toxins and adhesion factors (18). The *H. cinaedi* genome encodes a previously uncharacterized putative autotransporter protein. In this study, we named this uncharacterized protein *H. cinaedi* autotransporter protein A (HcaA) and investigated its contribution to *H. cinaedi* pathogenicity.

## RESULTS

### Structure and localization of HcaA

HcaA of *H. cinaedi* MRY 08-1234 [1,835 amino acids (aa), 197 kDa] is predicted to be composed of a signal peptide (1-27 aa), passenger domain (27-1425 aa), and β-barrel domain (1426-1825 aa, Fig. 1A). The β-barrel structure, characteristic of ATs, was highly supported based on the pLDDT score, indicating the high confidence of the model (Fig. 1A). Therefore, HcaA is predicted to be ATs. We constructed knockout mutants (Fig. 1B, top and middle) to identify the function of HcaA. *H. cinaedi* MRY08-1234 and MRY12-0027 strains were used as host strains, and *hcaA* was knocked out by replacing *hcaA* with *aphA* by homologous recombination (Fig. 1B, bottom). No *hcaA* complemented strains were constructed in this study, because no suitable plasmids for constructing complementation strains have been developed for *H. cinaedi*, and several trials have not succeeded in creating a *hcaA* complemented strain by inserting the *hcaA* gene on the chromosome. Therefore, we generated two mutants from strains with different backgrounds to confirm the effect of HcaA deletion. We had previously demonstrated the role of efflux pumps in *H. cinaedi* by creating several knockout strains in the same way (19). We believe that our approach (i.e., the use of multiple knockout strains) is suitable for investigating gene function. The knockout of *hcaA* was verified by polymerase chain reaction (PCR) amplification and immunoblotting using an anti-HcaA antibody (Fig. 1C and D). Immunoblot analysis revealed that HcaA was localized in the membrane fraction and was not secreted (Fig. 1D). Fluorescence immunostaining further confirmed the presence of HcaA on the bacterial surface (Fig. S1). No growth defects by the knockout of *hcaA* were observed (Fig. S2).

**Fig. 1.**
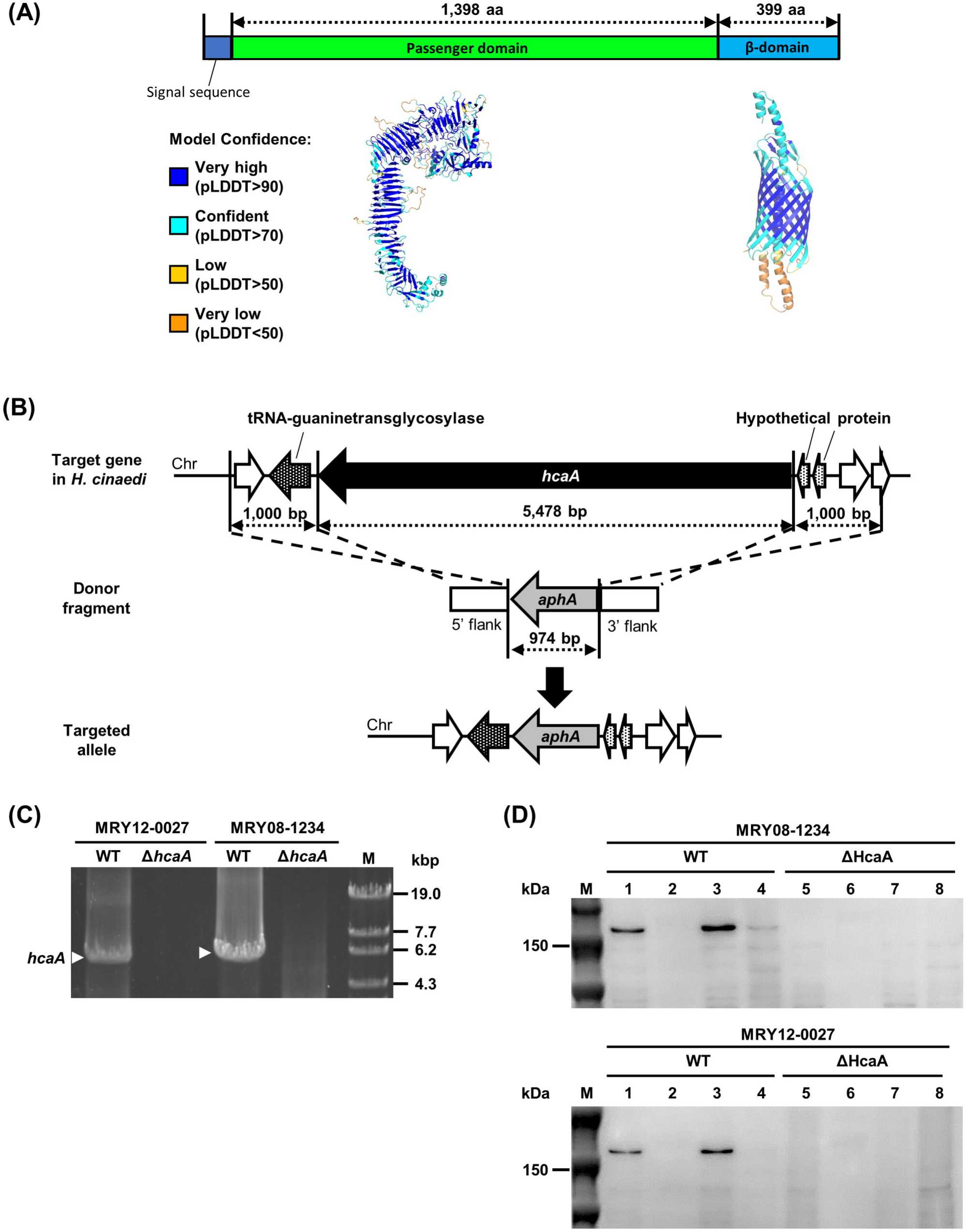
Construction of *hcaA*-knockout strains. (A) HcaA contains features of putative autotransporter proteins. The three-dimensional structure of HcaA predicted by AlphaFold2 is color-coded by pLDDT values, which indicate the confidence level of the prediction. (B) Schematic diagram of the knockout strategy for the *hcaA* gene. The *hcaA* locus on *H. cinaedi* MRY12-0027 and MRY08-1234 chromosomes, the donor fragment used to disrupt the *hcaA* gene locus, and the predicted *hcaA* gene locus after homologous recombination are shown. (C, D) Confirmation of *hcaA* gene knockout. (C) PCR using genomic DNA from wild-type (MRY08-1234 WT and MRY12-0027 WT) and HcaA-knockout strains (MRY08-1234 Δ*hcaA* and MRY12-0027 Δ*hcaA*) as templates. Lane 1, MRY12-0027 WT; Lane 2, MRY12-0027 Δ*hcaA*; Lane 3, MRY08-1234 WT; Lane 4, MRY08-1234 Δ*hcaA*; Lane 5, OneSTEP Marker 6 (λ/Sty I digest) (Nippon Gene Co., Ltd.). White arrows indicate amplicons of the *hcaA* gene (5,478 bp). Absence of the amplicons in HcaA-knockout strains was confirmed. (D) Immunoblot of HcaA. The concentration of the extracted protein was measured using the BCA protein assay. The samples were adjusted to a concentration of 0.5 mg/mL with PBS and separated by 5–20% sodium dodecyl sulfate-polyacrylamide gel electrophoresis. The top and bottom panels show results for MRY08-1234 and MRY12-0027, respectively. Lane M, Precision Plus Protein Dual Color Standards (Bio-Rad); Lane 1 and 5, whole cell lysate; Lane 2 and 6, culture supernatant fraction; Lane 3 and 7, membrane fraction; Lane 4 and 8, soluble fraction. The expected size of HcaA was 150 kDa, and a band corresponding to 150 kDa was observed in the WT (lanes 1 and 3) but not in the ΔHcaA strain (lanes 5 and 7), confirming the knockout and localization of HcaA.

### HcaA knockout reduced cytotoxicity and adherence of human colon epithelial cells *in vitro*

The effects of the HcaA knockout on cellular cytotoxicity were evaluated *in vitro* using a lactate dehydrogenase (LDH) assay. The human epithelial colorectal adenocarcinoma (Caco-2) cell line and human colon cancer (HT29) cell line were infected with *H. cinaedi* MRY08-1234 and MRY12-0027 strains. LDH production was significantly lower in cells infected with the HcaA-knockout strain than in those with the wild-type strain (Fig. 2A). In Caco-2 cells, the differences in LDH production were small, although yet significant differences were observed (*P* < 0.05). Therefore, HcaA may contribute to the cytotoxic effects of *H. cinaedi* infection on epithelial cells. HcaA knockout did not affect LDH production by mucus-secreting HT29-MTX cells (Fig. 2A), suggesting that the protective role of mucin in reducing the cytotoxicity of *H. cinaedi* was not affected by the knockout of HcaA. To investigate whether CDT is involved in the reduction of cytotoxicity, we analyzed its expression of *cdt* at the transcriptional level and observed no difference between the wild-type and HcaA-knockout strains (Figure S3).

**Fig. 2.**
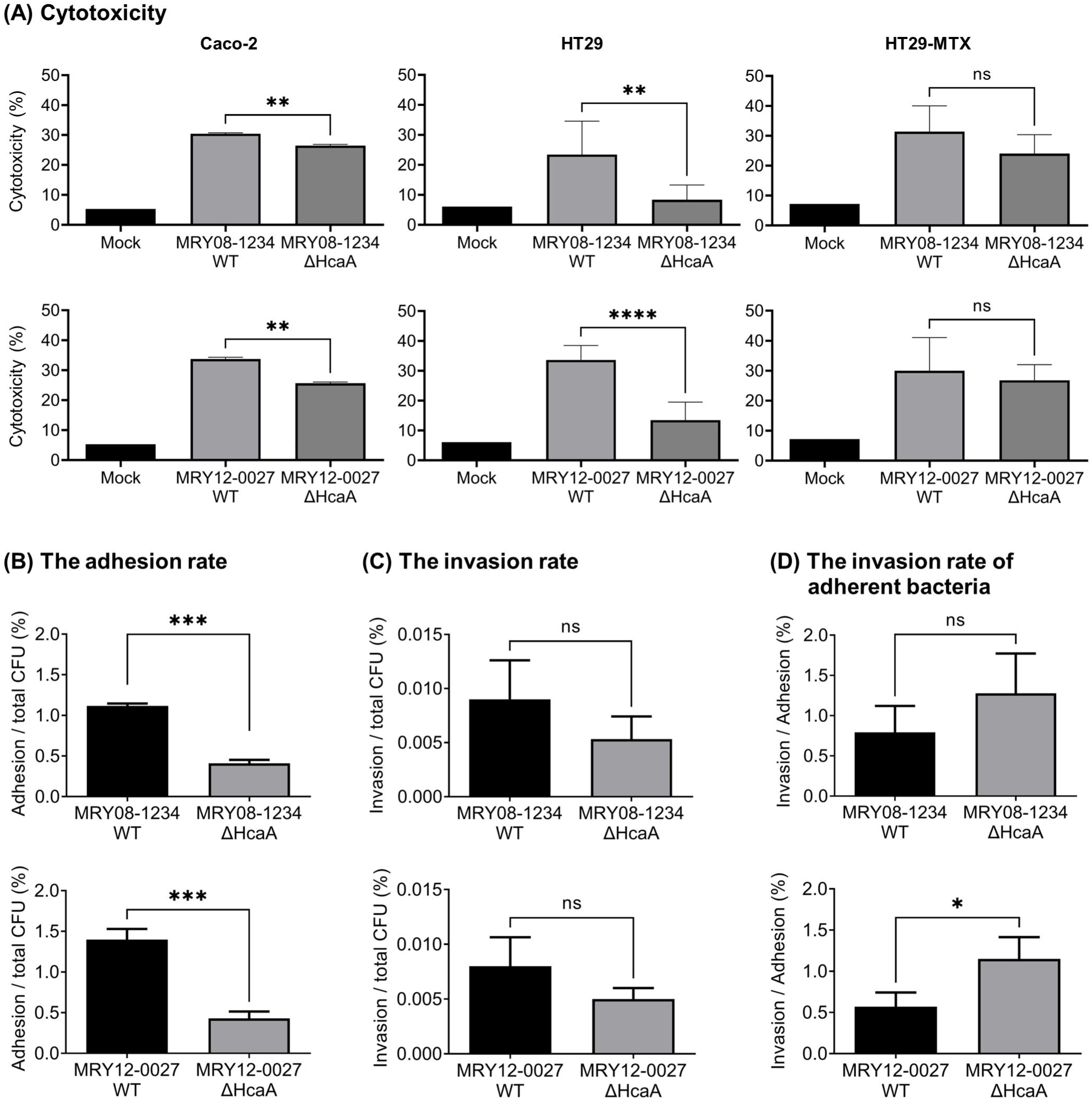
Knockout of HcaA reduced *H. cinaedi* cytotoxicity and adherence to cells. (A) LDH assay to evaluate cytotoxicity. Each cell was infected with wild-type (MRY08-1234 WT or MRY12-0027 WT) and HcaA-knockout (MRY08-1234 ΔHcaA or MRY12-0027 ΔHcaA) strains for 24 h (MOI 100). Cytotoxicity (%) was calculated using the method described in the instructions. (B), (C), and (D): Adhesion and invasion assays against Caco-2 cells. Caco-2 cells were infected with wild-type (MRY08-1234 WT or MRY12-0027 WT) and HcaA knockout (MRY08-1234 ΔHcaA or MRY12-0027 ΔHcaA) strains for 4 h (MOI 100). The adhesion rate (B) and invasion rate (C) were calculated by dividing the number of adherent bacterial cells and invading bacterial cells, respectively, by the total number of infected bacterial cells. The invasion rate of adherent bacteria (D) was calculated by dividing the number of invading bacteria cells by the number of adherent bacteria cells. Results are presented as the mean ± standard deviation (SD) of six independent experiments. *, *P*_<_0.05; **, *P* < 0.01; ***, *P* < 0.001 ****, *P*_<_0.0001; ns, not specified.

To determine whether HcaA contributes to cell adhesion and invasion, we performed a gentamicin protection assay, and compared the rate of adherent and invaded *H. cinaedi* cells between the wild-type and knockout strains. The adhesion rate of *H*. *cinaedi* MRY08-1234 to Caco-2 cells decreased significantly by HcaA knockout (Fig. 2B, upper panel). A significant reduction in the adhesion rate was also observed when *H. cinaedi* MRY12-0027 was used for gene knockout (Fig. 2B, bottom panel). However, it is important to note that this assay finding includes both bacteria adhered to the surface and those internalized within host cells. The invasion rates of MRY08-1234 and MRY12-0027 were also reduced by HcaA knockout; however, the differences were not significant (Fig. 2C). The invasion rate of adherent bacteria was higher in the HcaA-knockout strain and was significantly higher in MRY12-0027 (*P* <0.05, Fig. 2D).

### Adhesion was reduced in the recombinant mutant HcaA protein

HcaA contains an Arg-Gly-Asp (RGD) motif at amino acid residues 348-350. To determine the contribution of the RGD motif to adhesion, recombinant passenger regions of HcaA spanning 27-1,425 aa (out of total of 1,835 aa) of MRY08-1234 (WT HcaA) and mutant HcaA (with RGD motif replaced by RAD) were generated (Fig. 3A). Mutant HcaA showed a significant reduction in binding to U937 cells compared to that of WT HcaA (*P* < 0.05), indicating that the RGD motif contributes to binding to host cells (Fig. 3B). The predicted protein structure of HcaA of MRY08-1234 determined using AlphaFold2 indicated that the RGD motif is located on the surface of the protein, supporting the contribution of this motif to the adherence function of HcaA in *H. cinaedi* (Fig. 3C, left). The RGD motif was conserved in all *H. cinaedi* strains registered at the National Center for Biotechnology Information (NCBI), including clinical isolates (Fig. S4). HcaA homologs have been reported in *H. bilis*, *H. fennelliae*, *H. canicola*, and *H. canis*, all of which are closely related to *H. cinaedi* (20–22). The structural predictions for HcaA homologs determined using AlphaFold2 suggest that HcaA and its homologs have structural similarities in the passenger region; however, the RGD motif was present only in HcaA (Fig. 3C and D). In addition, the HcaA homologs are predicted to have a structure protruding from the surface, similar to the RGD motif of HcaA. However, the protruding structure did not contain the RGD motif (Fig. 3C).

**Fig. 3.**
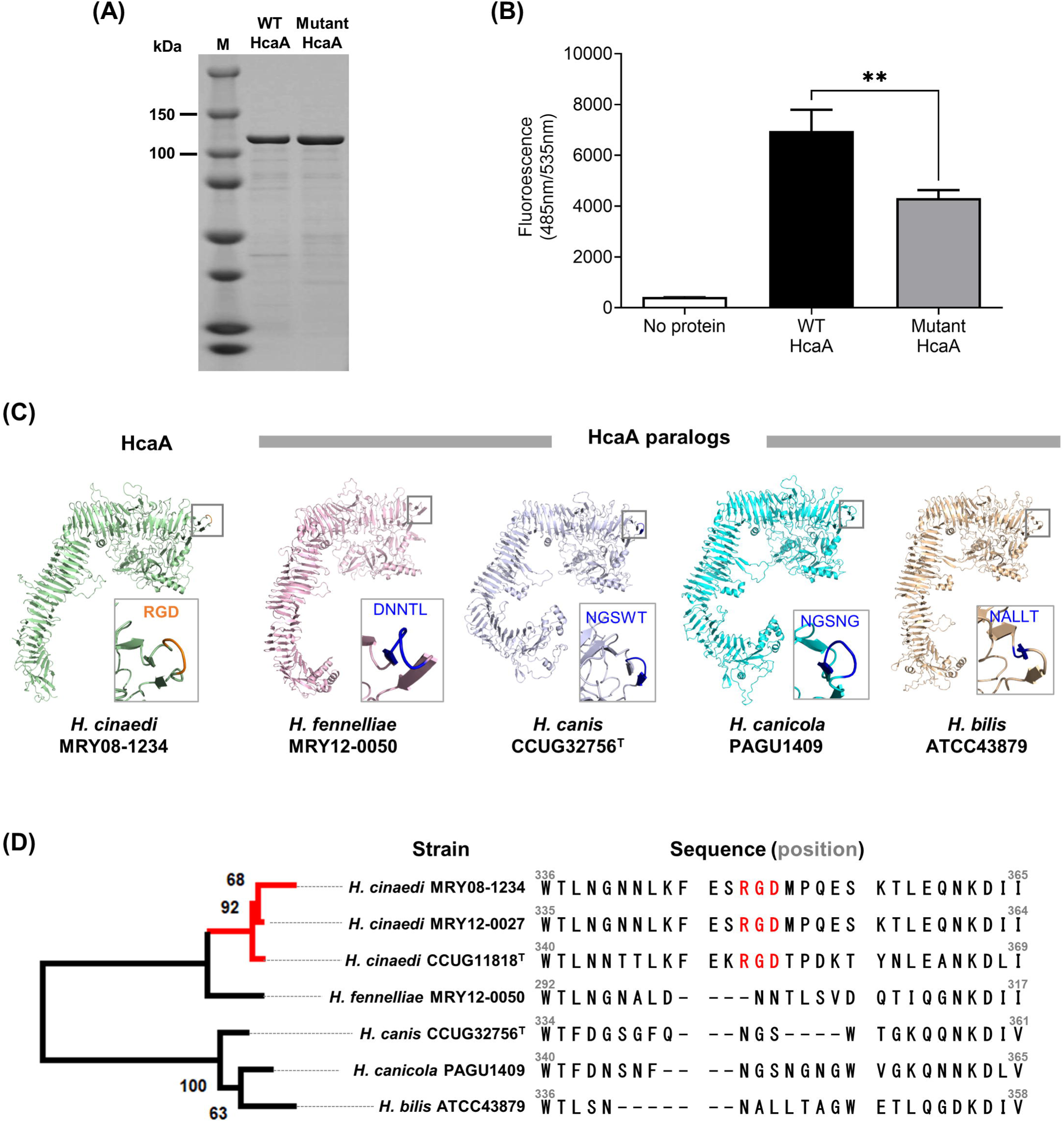
RGD motif on HcaA contributes to the adhesion of *H. cinaedi*. (A) SDS-PAGE of the purified WT HcaA and mutant HcaA, stained with Coomassie Brilliant Blue (CBB). HcaA (120 kDa) with the RGD motif (WT HcaA) and HcaA with glycine replaced with alanine in the RGD motif (Mutant HcaA) were expressed in *E. coli* and purified. Proteins were adjusted to 1 mg/mL and applied. (B) Adhesion of purified WT HcaA (with the RGD motif) and Mutant HcaA (with the RAD motif) to U937 cells. Fluorescence-labeled U937 cells attached to HcaA were compared by measuring the fluorescence at 490 nm. Means and SD reflect values of six per group. **, *P*_<_0.01. (C) Three-dimensional structure of HcaA and HcaA homologues predicted by AlphaFold2. Of the five models generated, the top-ranked model was used in this study. The predicted structure was visualized using PyMOL Molecular Graphics System Ver. 2.2.5. HcaA homologs from *H. bilis*, *H. fennelliae*, *H. canicola*, and *H. canis* were analyzed and compared. The RGD motif is located at the protrusion in HcaA (within the square). HcaA homologs had a structure protruding from the surface, like HcaA; however, the protruding structure did not contain the RGD motif. (D) Phylogenetic tree based on the alignment of the passenger domain of HcaA and HcaA homologs. The alignment was generated using ClustalW ver. 2.1. The maximum-likelihood tree was constructed using RAxML (https://github.com/amkozlov/raxml-ng) and visualized using MEGA-X v 10.2.6. A comparison of the amino acid sequence of HcaA showed that the RGD motif was present only in HcaA from *H. cinaedi* (red text).

### Effects of HcaA on the establishment of *H. cinaedi* colonization in mice

To clarify the role of HcaA in *H. cinaedi* infection *in vivo*, C57BL/6 female mice were orally infected with wild-type and HcaA-knockout *H. cinaedi* strains. The recovery of *H. cinaedi* from the colon, cecum, cecal contents, biceps femoris muscle, and trapezius muscle was analyzed at 7, 14, and 28 days post-infection (dpi). Because *H. cinaedi* is known to cause cellulitis and infected aortic aneurysm (23), muscle samples were also collected in addition to the intestinal tract samples. Animal experiments were performed twice, yielding consistent results. Results of the first experiment, including only the MRY08-1234 strain, are shown in Fig. S5. The results of the second experiment, including both MRY08-1234 and MRY12-0027 strains, are shown in Figs. 4 and S6. To confirm colonization, reactivity to *H. cinaedi* was measured in mouse serum. An increase in reactivity against whole cells of *H. cinaedi* was observed in both groups of infected mice. Reactivity to *H. cinaedi* tended to be higher in the group of mice infected with the wild-type strain than in the mice infected with the knockout strain (Fig. 4A). In the MRY08-1234 strain, the reactivity at 14 dpi was significantly higher than that in the knockout strain (*P* < 0.01). Using the culture method, *H. cinaedi* was recovered from tissues other than cecal contents in both groups of mice infected with the wild-type and HcaA-knockout strains (Table 1). The amount of *H. cinaedi* was highest in the cecum from mice infected with the wild-type strains (MRY08-1234 WT and MRY12-0027 WT) at 7 dpi. The recovery rate of *H. cinaedi* from the cecum was lower in mice infected with the HcaA-knockout strain (0%) than in mice infected with the wild-type strain (MRY08-1234 WT, 2/6, 33.3%; MRY12-0027 WT, 2/5, 40.0%, *P* = 0.12). At 14 dpi, there were lower rates of colonization in the cecum of mice infected with the HcaA-knockout strain (MRY08-1234 KO, 0/5, 0%; MRY12-0027 KO, 0/5, 0%) strain than in mice infected with the wild-type strain (MRY08-1234, 4/5, 80.0%; MRY12-0027, 2/5, 40.0%, *P* = 0.06). In MRY08-1234, there was a trend towards a lower rate of colonization in the colon of mice infected with the HcaA-knockout strain than with the wild-type strain at 14 dpi. Similarly, in MRY12-0027, there were also lower rates of colonization in mice infected with the HcaA-knockout strain (MRY12-0027 ΔHcaA, 0/5, 0%) than in mice infected with the wild-type strain (MRY12-0027 WT, 1/5, 20%). *H. cinaedi* was not isolated from the cecal contents. For the cecum and colon, in which bacteria were frequently detected by the culture method, mRNA was extracted and bacteria were detected by quantitative real-time PCR (qPCR). The frequency of positive samples was higher in the group of mice infected with the wild-type strain than in mice infected with the knockout strain (Fig. S6). Therefore, a combined analysis of the results of the culture and qPCR assays showed that the frequency of positive detection was higher for mice infected with the wild-type strain than in mice infected with the HcaA-knockout (Fig. 4B). A reduction in the number of HcaA-knockout bacterial cells was also observed by an immunofluorescence analysis of the colon infected with *H. cinaedi* using an anti*-H. cinaedi* antibody (Figs. 4C and S5). Regardless of the presence or absence of HcaA and the duration of infection, *H. cinaedi* was observed in the mucosal epithelium. However, in samples from mice infected with the HcaA-knockout strain, the number of bacteria tended to be lower compared to samples from mice infected with the wild-type strain.

**Fig. 4.**
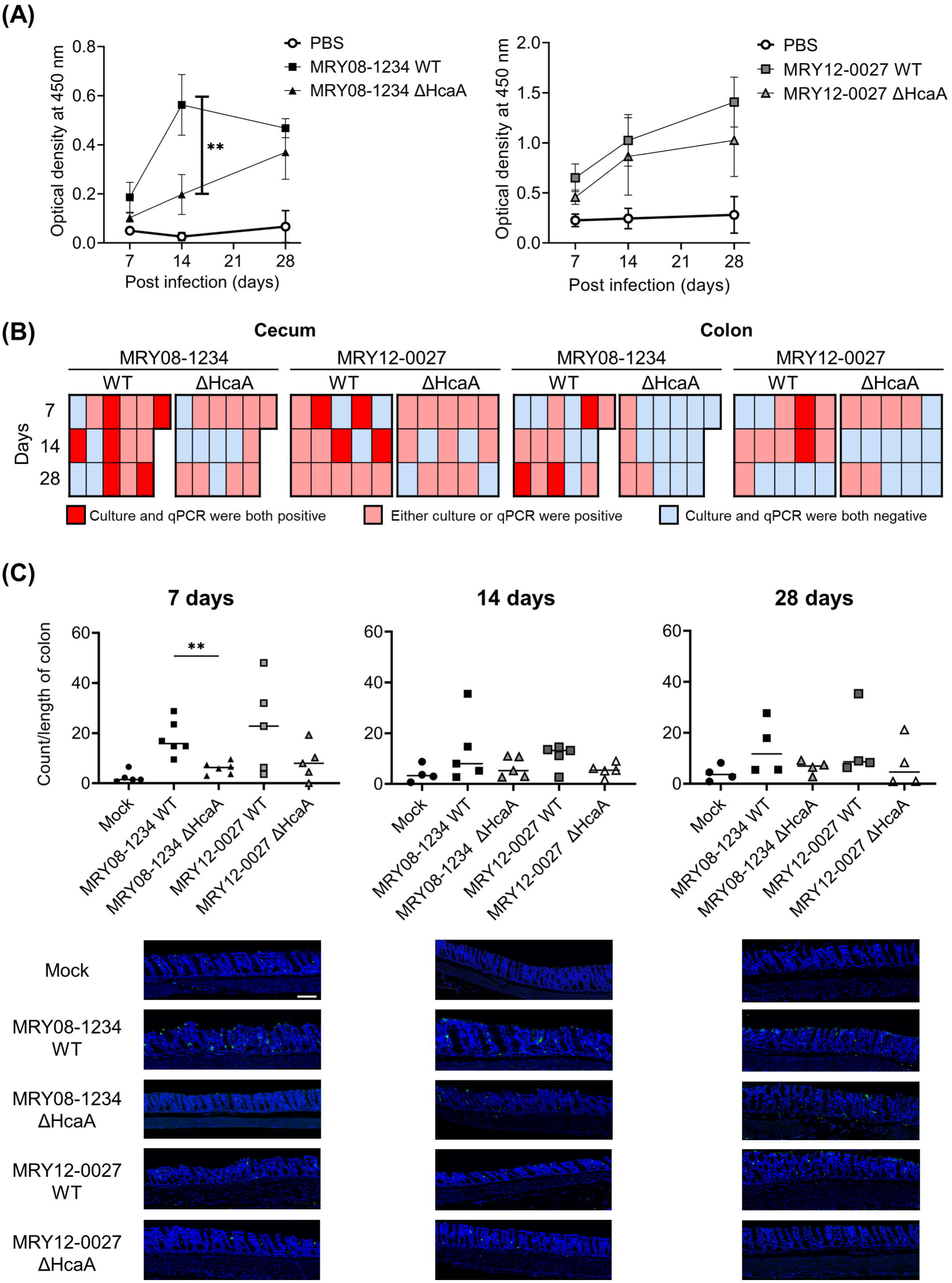
Knockout of HcaA reduced *H. cinaedi* colonization in mice. (A) Reactivity to *H. cinaedi* in infected mice, as determined by ELISA. Sera from mice at 7, 14, and 28 days after inoculation were used. Squares and triangles indicate serum from the group of infected wild-type (black square, MRY08-1234 WT; gray square, MRY12-0027 WT) or HcaA knockout (black triangle, MRY08-1234 ΔHcaA; gray triangle, MRY12-0027 ΔHcaA) strains, respectively. Means and SD reflect values from five of six mice per group. **, *P*_<_0.01. (B) Heatmap combining the results of the culture method and qPCR assays. Columns indicate each mouse and rows indicate the duration of infection. Positive results for both culture method and qPCR assays are shown in red, positive results for either are shown in pink, and negative results for both are shown in light blue. (C) Quantitative bacteria count per length of colon and representative images of the colon of *H. cinaedi*-infected mice. Adhesion of *H. cinaedi* to the colon was quantified using ImageJ ver. 1.53t, by counting the number of *H. cinaedi* in images taken with a 20× objective lens. Results are expressed as the number of *H. cinaedi* per length of colon. Means and SD reflect the values of five or six mice per group. **, *P*_<_0.01. Green, *H. cinaedi*; blue, nucleus. Scale bar 100 um.

**Table 1.** Number of *H. cinaedi* isolated from tissues of *H. cinaedi*-infected mice.

| Tissue<sup>#</sup> | Bacterial count (CFU g<sup>-1</sup>) by infection duration |  |  |  |  |  |  |  |  |  |  |  |
| --- | --- | --- | --- | --- | --- | --- | --- | --- | --- | --- | --- | --- |
|  | MRY08-1234 WT |  |  | MRY08-1234 KO |  |  | MRY12-0027 WT |  |  | MRY12-0027 KO |  |  |
|  | 7 d | 14 d | 28 d | 7 d | 14 d | 28 d | 7 d | 14 d | 28 d | 7 d | 14 d | 28 d |
| Cecum |  |  |  |  |  |  |  |  |  |  |  |  |
| 1 | 0 | 2.0×10<sup>3</sup> | 0 | 0 | 0 | 0 | 0 | 0 | 0 | 0 | 0 | 0 |
| 2 | 0 | 0 | 0 | 0 | 0 | 0 | 9.5×10<sup>5</sup> | 0 | 0 | 0 | 0 | 1.5×10<sup>3</sup> |
| 3 | 1.5×10<sup>5</sup> | 1.0×10<sup>3</sup> | 1.0×10<sup>3</sup> | 0 | 0 | 0 | 0 | 1.0×10<sup>3</sup> | 0 | 0 | 0 | 0 |
| 4 | 0 | 1.0×10<sup>3</sup> | 0 | 0 | 0 | 0 | 1.5×10<sup>5</sup> | 0 | 0 | 0 | 0 | 0 |
| 5 | 0 | 1.5×10<sup>5</sup> | 2.0×10<sup>3</sup> | 0 | 0 | 1.5×10<sup>3</sup> | 0 | 2.0×10<sup>3</sup> | 0 | 0 | 0 | 0 |
| 6 | 9.5×10<sup>5</sup> | - | - | 0 | - | - | - | - | - | - | - | - |
| Colon |  |  |  |  |  |  |  |  |  |  |  |  |
| 1 | 0 | 0 | 2.5×10<sup>3</sup> | 0 | 0 | 1.0×10<sup>3</sup> | 0 | 0 | 9.0×10<sup>3</sup> | 0 | 0 | 1.0×10<sup>3</sup> |
| 2 | 0 | 2.5×10<sup>3</sup> | 2.0×10<sup>3</sup> | 0 | 0 | 2.0×10<sup>2</sup> | 0 | 0 | 0 | 0 | 0 | 2.0×10<sup>2</sup> |
| 3 | 3.0×10<sup>3</sup> | 0 | 2.0×10<sup>3</sup> | 0 | 0 | 0 | 5.0×10<sup>2</sup> | 0 | 0 | 0 | 0 | 0 |
| 4 | 0 | 0 | 0 | 0 | 0 | 0 | 3.0×10<sup>3</sup> | 2.5×10<sup>3</sup> | 0 | 0 | 0 | 0 |
| 5 | 5.0×10<sup>2</sup> | 0 | 9.5×10<sup>3</sup> | 0 | 0 | 0 | 0 | 0 | 0 | 0 | 0 | 0 |
| 6 | 0 | - | - | 0 | - | - | - | - | - | - | - | - |

Table 1. (Continued.)
| Bacterial count (CFU g<sup>-1</sup>) by infection duration |  |  |  |  |  |  |  |  |  |  |  |  |
| --- | --- | --- | --- | --- | --- | --- | --- | --- | --- | --- | --- | --- |
| Tissue | MRY08-1234 WT |  |  | MRY08-1234 KO |  |  | MRY12-0027 WT |  |  | MRY12-0027 KO |  |  |
|  | 7 d | 14 d | 28 d | 7 d | 14 d | 28 d | 7 d | 14 d | 28 d | 7 d | 14 d | 28 d |
| Biceps femoris muscle |  |  |  |  |  |  |  |  |  |  |  |  |
| 1 | 0 | 0 | 0 | 0 | 0 | 0 | 0 | 0 | 0 | 0 | 0 | 0 |
| 2 | 0 | 0 | 2.0×10<sup>2</sup> | 0 | 5.0×10<sup>2</sup> | 1.0×10<sup>1</sup> | 0 | 0 | 0 | 0 | 0 | 0 |
| 3 | 0 | 0 | 0 | 0 | 0 | 0 | 0 | 0 | 0 | 0 | 0 | 0 |
| 4 | 0 | 0 | 0 | 0 | 0 | 0 | 0 | 0 | 0 | 0 | 0 | 0 |
| 5 | 0 | 0 | 0 | 0 | 0 | 0 | 0 | 0 | 0 | 0 | 0 | 0 |
| 6 | 0 | - | - | 0 | - | - | - | - | - | - | - | - |
| Trapezius muscle |  |  |  |  |  |  |  |  |  |  |  |  |
| 1 | 0 | 0 | 0 | 0 | 0 | 0 | 0 | 0 | 0 | 0 | 0 | 0 |
| 2 | 0 | 0 | 0 | 0 | 0 | 0 | 0 | 0 | 0 | 0 | 0 | 0 |
| 3 | 0 | 5.0×10<sup>2</sup> | 0 | 0 | 0 | 0 | 0 | 5.0×10<sup>2</sup> | 0 | 0 | 0 | 0 |
| 4 | 0 | 0 | 0 | 0 | 0 | 0 | 0 | 0 | 0 | 0 | 0 | 0 |
| 5 | 0 | 0 | 0 | 0 | 0 | 0 | 0 | 0 | 0 | 0 | 0 | 0 |
| 6 | 0 | - | - | 0 | - | - | - | - | - | - | - | - |

<sup>#</sup>Cecal contents of all mice were cultured, and *H. cinaedi* was not isolated. WT, wild type; KO, HcaA-knockout

## DISCUSSION

Enterohepatic *Helicobacter* species are increasingly recognized as critical pathogens in mammals. Among them, *H. cinaedi* causes bacteremia, cellulitis, and infected aortic aneurysms in humans (2, 4, 24). Although *H. cinaedi* infections are rarely reported, cases have been increasing in Japan since 2003 (8, 11, 24). However, little is known about the virulence factors of *H. cinaedi*. ATs have various functions that contribute to the establishment of infections by pathogenic gram-negative bacteria. In this study, we found that HcaA, a putative AT, contributes to adhesion to host cells and to the colonization of *H. cinaedi*. This is the first report to elucidate the role of ATs in enterohepatic *Helicobacter* species.

T5SS proteins are classified into subgroups labelled Va–Ve; however, this classification is based on protein architecture and does not reflect the functions of the secreted passengers. VacA produced by *H. pylori* is classified as a type Va AT; it binds to host cells and is internalized, causing vacuolation characterized by the accumulation of large vesicles (25). Serine protease autotransporters of *Enterobacteriaceae* (SPATEs) produced by *Enterobacteriaceae* are also classified as type Va ATs (26). SPATEs exhibit protease activity to cleave host proteins. HcaA is structurally similar to EspP, SPATEs in *Escherichia coli*. We did not detect protease activity against casein and pepsin, the major substrates for SPATEs (Fig. S7a and b). Furthermore, no significant differences in CDT expression were observed in the presence or absence of HcaA (Fig. S3). In contrast, the number of bacterial cells adhering to human epithelial cells and the level of cytotoxicity were significantly reduced by HcaA knockout. These findings suggest that the reduced cytotoxicity observed in the HcaA-knockout strain can be explained by the decrease in the number of adherent *H. cinaedi* cells, rather than a direct cytotoxic effect of HcaA itself.

HcaA contains the RGD motif in its passenger region. The RGD motif acts as a fibronectin cell adhesion site. It plays a role in adhesion to host epithelial cells that express receptors of the RGD motif, such as integrins (27). Cell adhesion experiments using purified WT HcaA with the RGD motif and mutant HcaA with the RAD motif showed that the adhesion ability decreased significantly when RGD was replaced by RAD. Structural prediction indicated that the RGD motif is located on the surface of HcaA at an externally protruding site. The RGD motif is known to bind to integrins, which are thought to mediate the adhesion of HcaA to cells. Our results strongly suggest that the RGD motif of HcaA plays a role in cell adhesion. However, the adhesion ability of HcaA was not completely diminished by replacing RGD with RAD. Therefore, HcaA may function as an adhesin independent of the RGD motif, and the presence of the RGD motif enhances its ability to adhere to cells. HcaA homologs in other enterohepatic *Helicobacter* species, such as *H. bilis*, *H. canis*, *H. fennelliae,* and *H. canicola,* lacked the RGD motif (Fig. 3). In addition, the RGD motif of HcaA was highly conserved among *H. cinaedi* strains, regardless of strain origin (Fig. S4). *H. canicola* is genetically most closely related to *H. cinaedi* (22). However, a phylogenetic analysis indicated that HcaA is most similar to the HcaA homolog of *H. fennelliae* and not to that of *H. canicola*. *H. fennelliae* is the second most common enterohepatic *Helicobacter* species causing invasive infections in humans. The structural similarity between HcaA and HcaA homologs of *H. fennelliae* suggests that HcaA plays a critical role in the pathogenesis of enterohepatic *Helicobacter* species in humans. The results of comparative genomic analyses indicate that *H. cinaedi* is a human-adapted lineage, unlike *H. canicola* (22). The acquisition of the RGD motif of HcaA during the evolution of *H. cinaedi* may have contributed to the adaptation of *H. cinaedi* to humans, resulting in a higher isolation rate of *H. cinaedi* from humans compared to other enterohepatic *Helicobacter* species. Further investigation is needed to identify the HcaA-binding target in the host cell.

The mouse infection experiment also supported the results of *in vitro* analyses, showing that HcaA knockout reduced the number of infected bacterial cells in the cecum and colon of *H. cinaedi*-infected mice. In particular, the HcaA-knockout strain had significantly fewer bacterial cells in culture than that in the wild-type strain at 7 dpi. Therefore, HcaA plays an important role in the established phase of *H. cinaedi* infection. A similar effect was observed for FaaA (flagella-associated autotransporter A), an AT possessed by *H. pylori* contributing to gastric colonization, and an FaaA deficiency has been reported to significantly reduce the number of colonized bacteria in the early stages of infection (28).

ATs have various functions. For example, ImaA, another ATs in *H. pylori,* is essential for host colonization and modulates the host inflammatory response via the inflammatory cytokines IL-8 and TNF-α (29). In addition, EspC, SPATEs, degrades various host proteins, such as hemoglobin and mucin, and also to prevent cytotoxicity during the initial stage of bacterial infection by regulating pore formation (26, 30). Thus, similar to other ATs, HcaA can also have multiple functions related to the establishment of chronic *H. cinaedi* infections in humans. Further analyses are needed to elucidate additional functions of HcaA beyond its role in cell adhesion.

This study identified a novel virulence factor of *H. cinaedi*, HcaA, as an adhesin protein. HcaA was shown to play an important role in establishing *H. cinaedi* infection both *in vitro* and *in vivo*. On the other hand, adhesion was also observed in the HcaA-knockout strain, although it was reduced compared to that of the wild-type strain, suggesting that factors other than HcaA contribute to host adhesion. Further studies of HcaA and other factors are needed to understand the interaction between *H. cinaedi* and host cells.

## MATERIALS AND METHODS

### Condition of bacterial strains and host cells

*H. cinaedi* strains MRY08-1234 (MLST: ST10) and MRY12-0027 (MLST: ST15) isolated from the blood of patients with bacteremia in Japan were used throughout the study. The *H. cinaedi* strains were cultured as described previously (19). Briefly, strains were subcultured on Brucella agar (Becton, Dickinson, Franklin Lakes, NJ, USA) with 5% horse blood at 37 °C under microaerobic conditions with hydrogen generated by the gas exchange method using an anaerobic gas mixture (H_2_, 10%; CO_2_, 10%; and N_2_, 80%). For liquid culture, bacteria were suspended in Brucella broth (Becton, Dickinson) with 10% fetal bovine serum (FBS; NICHIREI), and the culture was incubated at 37 °C under microaerobic conditions with hydrogen.

Human epithelial colorectal adenocarcinoma cells (Caco-2), a human myelomonocyte cell line (U937), a human colon cancer cell line (HT29), and mucus-secreting HT29 cells (HT29-MX) were used in this study. Caco-2 cells were maintained in Eagle’s minimum essential medium (EMEM, Sigma Aldrich, St. Louis, MO, USA) containing 2 mM L-glutamine, 1 mM sodium pyruvate, 1.5 mg/L sodium bicarbonate, 10 mg/L phenol red, 10% heat-inactivated (56 °C, 30 min) FBS, 1% non-essential amino acids. U937, HT29, and HT29-MX cells were maintained in RPMI 1640 (Sigma Aldrich) with 10% heat-inactivated FBS. All media contained an antimicrobial cocktail (penicillin-streptomycin), except for the *H. cinaedi* infection experiments. All cells were incubated in a 5% CO_2_, humidified atmosphere at 37 °C, in a stationary manner.

### Generation of *hcaA* deletion mutants and antibodies to HcaA

Knockout mutants were generated according to previously described methods (19). Briefly, 1,000 base pairs up– and downstream of *hcaA* were PCR-amplified. The primers contained a linker sequence overlapping with *aphA*. To obtain the construct of *aphA* sandwiched by the target gene sequence, three purified PCR products, up– and downstream of *hcaA* and *aphA*, were mixed and amplified. The donor fragment was electroporated into *H. cinaedi*. The transformed *H. cinaedi* was plated on agar plates containing 10 µg/mL kanamycin and 3% agar. After incubation, the growing colonies were selected as knockout mutants. The knockout of *hcaA* was confirmed by PCR and an immunoblot analysis. Primers used for cloning and PCR are listed in Table S1. For the immunoblot analysis, the whole bacterial cell lysates, culture supernatant, soluble fraction, and membrane fraction were prepared as follows. The bacterial culture was incubated until an optical density at 600 nm (OD600) of 0.4, and the supernatant was collected by centrifugation (8,600 ×*g*, 10 min, 4 °C). Then, the insoluble and the soluble fractions were prepared by extracting proteins from the pellet using the EzBactYeast Crusher (ATTO Corp., Tokyo, Japan) and by centrifugation. The insoluble fraction was used as the membrane fraction. These samples were separated and transferred to Immobilon^TM^ –FL PVDF (pore size, 0.45 um; Millipore Inc., Burlington, MA, USA) using a semidry electroblotting system. An anti-HcaA antibody was obtained by immunizing rabbits with a KLH-conjugated peptide (RGDMPQESKTLEQN corresponding to positions 348–361 of HcaA in MRY08-1234), followed by affinity purification against the peptide. The resulting rabbit polyclonal HcaA antibody was used as the primary antibody for the immunoblot analysis (1:1,000), and peroxidase-conjugative AffiniPure donkey anti-mouse IgG (H+L) (Jackson ImmunoResearch Inc., 1:50,000) was used as the secondary antibody. The bands were visualized using ECL Select (Amersham Biosciences, Amersham, UK) and Amersham^TM^ ImageQuant 800 (Amersham Biosciences).

### Cell infection experiments

Strains were grown in Brucella broth with 10% FBS at 37 °C under microaerobic conditions. After incubation, the bacterial culture was inoculated into 10 mL of fresh broth, and the culture was incubated until an OD600 of 0.4. Bacterial cells were harvested by centrifugation (3,350 × *g*, 10 min, 25 °C) and resuspended in the appropriate medium for each cell. Infections were performed at a multiplicity of infection (MOI) of 100. Immediately after the addition of bacteria, the plates were centrifuged at 300 ×*g* for 5 min at 25 °C, followed by incubation in 5% CO_2_ humidified atmosphere, at 37 °C for 4 h.

Cytotoxicity was evaluated by measuring lactate dehydrogenase using the Cytotoxicity LDH Assay Kit-WST (Dojindo Laboratories, Tokyo, Japan). The assay was performed according to the manufacturer’s instructions.

Cell adhesion and invasion ability were evaluated using a gentamicin assay. To evaluate adhesion ability, cells were liberated by trypsinization, seeded into a 24-well tissue culture-treated polystyrene flat-bottom microtiter plate (FALCON), and incubated to confluence. Caco-2 cells were infected by adding 1 mL of bacterial suspension (MOI 100) directly to each well. After 4 h of incubation, the un-bound bacteria were removed by washing with pre-warmed PBS. To evaluate invasive ability, *H. cinaedi* adherent to Caco-2 cells was incubated in EMEM with gentamicin at 250 µg/mL. After 1.5 h, the infected cells were washed with pre-warmed PBS. For both assays, host cells were washed with PBS containing 0.1% saponin (FUJIFILM Wako Pure Chemical Co., Osaka, Japan). Serial dilutions of the suspension were plated on Brucella agar containing 5% horse blood and 3% agar. After incubation, colonies were counted, and the adhesion rate (%), invasion rate (%), and the invasion rate of adherent bacteria (%) were calculated.

### Purification of HcaA

To produce recombinant HcaA, we constructed expression vectors; *hcaA* was amplified by PCR and introduced into pCold™ I DNA (TaKaRa, Shiga, Japan) using the In-Fusion® HD Cloning Kit. gDNA from *H. cinaedi* strain MRY08-1234 was used to amplify the passenger region of HcaA (27-1,425/1,835 aa). The expression vector containing the passenger region of HcaA was transferred into *Escherichia coli* BL21 (DE3) pLysS cells and plated on LB agar plates containing 100 µg/mL ampicillin. The growing colonies were inoculated into 100 mL of LB broth containing 100 µg/mL ampicillin and incubated at 37 °C with shaking at 200 rpm for 3 h. The cells were then transferred to 900 mL of fresh medium and grown in a 2 L flask at 37 °C until OD600 of 0.6. Protein expression was induced by cooling in ice water for 1 h and adding 400 µL of 1 mM isopropyl β-D-thiogalactopyranoside. Recombinant HcaA was expressed as a fusion protein containing a histidine tag. The cell cultures were further incubated at 15 °C overnight. Bacteria were harvested by centrifugation at 5,000 ×*g* for 10 min at 4 °C and resuspended in 20 mM Tris-HCl (Nacalai, Kyoto, Japan), 150 mM NaCl (FUJIFILM Wako), and 40 mM imidazole (FUJIFILM Wako) at pH 7.6 with cOmplete™ Protease Inhibitor Cocktail (Roche Molecular Systems). The suspended solution was sonicated on ice, and the supernatant was collected by centrifugation at 5,000 ×*g* for 45 min at 4 °C. The recombinant protein was purified in two steps using an affinity column (HisTrap FF crude, Cytiva), followed by size exclusion chromatography (Superdex™ 200 Increase 10/300 GL, Cytiva). To replace the RGD motif of the HcaA protein with RAD, site-directed mutagenesis of the HcaA expression vector was performed using PrimeSTAR^®^ GXL DNA Polymerase. The primers used to generate expression vectors are listed in Table S1.

### Adhesion assay of HcaA to U937 cells

Purified HcaA and mutant HcaA were prepared at a concentration of 50 µg/mL in PBS, dispensed in 100 µL aliquots to 96-well plates, and fixed to the bottom of the plates by incubation overnight at 4 °C. The wells were then washed with PBS and blocked by adding 2% BSA in PBS and incubated again overnight at 4 °C. U937 cells were incubated until confluence and then diluted with MEM to a concentration of 1 × 10^5^ cells/mL. The use of U937 cells in this assay was based on previous analyses of their activity in relation to *H. pylori* (31). Fluorescein (FUJIFILM Wako) was added to the cell suspension at a final concentration of 1 mg/mL and incubated at 37 °C for 30 min. After washing, MEM containing 2 mM Ca and 2 mM Mg was added and allowed to react for 30 min on ice in the dark. The cell suspension, diluted to a concentration of 5 × 10^5^ /mL, was then added to HcaA-coated plates and incubated at 37 °C for 1 h. After washing and replacement with phenol red-free MEM medium (Sigma), fluorescence was measured, and the number of cells bound to purified proteins was observed.

### Sequence alignment, phylogenetic analysis, and structure prediction of HcaA and HcaA homologs

Amino acid sequences of HcaA homologs in other enterohepatic *Helicobacter* species were obtained from the whole genome sequences of *H. bilis* ATCC43879 (Accession No. KI392040), and *H. fennelliae* MRY12-0050 (Accession No. BSAD00000000), and *H. canicola* PAGU1409 (Accession no. DRR092854) and *H. canis* CCUG32756 (Accession No. GCF008693005) were obtained from the National Center for Biotechnology Information (NCBI). The passenger regions of the amino acid sequences of the HcaA and HcaA homologs were aligned using ClustalW ver. 2.1 (32), and visualized using CLC Genomics Workbench Version 22.0.1 (QIAGEN). A phylogenetic tree based on the aligned amino acid sequences of the passenger domain of HcaA and HcaA homologs was constructed using RAxML-NG v1.1.0 (https://github.com/amkozlov/raxml-ng) and visualized using MEGA X v 10.2.6 (33).

The prediction of the structures of HcaA and HcaA homologs were performed using AlphaFold2 Ver.2.2.2 on the supercomputer system of the Institute for Chemical Research, Kyoto University, with a full dBs preset.

### Animal experiments and infection

Animal experiments were performed twice; the results of the first animal experiment are shown in the Supplementary Appendix, and the results of the second experiment are reported in the manuscript. Six-week-old female C57BL/6 mice (Japan SLC, Inc., Shizuoka, Japan) maintained under specific pathogen-free conditions were used in this study. Mice were fed a standard rodent chow, given ad libitum access to water, housed in microisolator cages, and isolated from specific pathogens.

Strains were suspended in PBS at 1 × 10^9^ cells/mL. The mice were then orally administered 0.2 mL of each bacterial suspension through a 1 mL syringe attached to an oral probe every other day for a total of three times. The same volume of PBS was administered to mice as a negative control. There were 14 mice in the control group, 15 mice in the wild-type strain infection group, and 15 mice in the HcaA-knockout strain infection group.

On days 7, 14, and 28 after inoculation, five or six mice from each group were sacrificed for analyses. The blood, cecal contents, colon, biceps femoris muscle, and trapezius muscle were collected aseptically for further analysis. All animal experiments and the study protocol were approved by the Committee for Animal Experimentation of the National Institute of Infectious Diseases (approval number 121126).

## ELISA

To measure reactivity to *H. cinaedi* in mouse serum, an enzyme-linked immunosorbent assay (ELISA) was performed as described previously (34). To prepare serum, whole blood collected at autopsy was incubated at 4 °C and then centrifuged at 2,000 ×*g* for 5 min at 4 °C. The supernatant was collected, mixed with glycerol (50% final concentration), and designated as serum. To prepare whole-cell lysates, *H. cinaedi* MRY08-1234 cells were disrupted by sonication. Each well of Nunc-Immuno 96-well microtiter plates (No. 439454; Thermo Fisher Scientific, Waltham, MA, USA) was coated with of whole-cell lysates (4 ug/mL in 0.1 M carbonate/bicarbonate buffer, pH 9.4). After overnight incubation at 4 °C, the plate was washed with PBS containing 0.05% (vol/vol) Tween 20 (PBS-T), and the wells were saturated with blocking buffer (PBS containing 1% BSA) and incubated for 1 h at 37 °C while shaking at 500 rpm. After washing with PBS-T, the wells were filled with serum samples, which were diluted 100-fold with a blocking buffer. The dilution of serum was determined by preliminary experiments (Fig. S8). Plates were incubated for 1 h at 37 °C with shaking at 500 rpm. Horseradish peroxidase-conjugated goat anti-mouse IgG secondary antibody (Jackson, Bar Harbor, ME, USA) diluted 5,000-fold in blocking buffer was used. Absorbance was measured at 450 nm using a Multiskan FC microplate reader (Thermo Scientific).

### Isolation and detection of *H. cinaedi* from infected mice

Culture and PCR were used to assess *H. cinaedi* infection in each organ. For the cultivation of *H. cinaedi*, agar plates containing Skirrow (Oxoid), 5% horse blood, 3% agar, and 4 µg/mL ciprofloxacin (FUJIFILM Wako) were inoculated with the cecal contents, colon, biceps femoris muscle and trapezius muscle that had been homogenized in Brucella broth. MRY08-1234 is resistant to ciprofloxacin; accordingly, ciprofloxacin was added to eliminate commensal intestinal bacteria in mice and isolate only *H. cinaedi*. Plates were incubated at 37 °C under microaerobic conditions. The growth of the colonies was quantified and confirmed via detection of *cdt* using PCR to determine whether they were colonies of *H. cinaedi*. Data for MRY08-1234 WT and MRY08-1234 KO at 7 dpi were collected from six mice in each group, while data for other strains were collected from five mice in each group. We also evaluated total RNA of *H. cinaedi*. Samples were homogenized with 1 mL of Isogen (Nippon Gene, Toyama, Japan). The homogenates were supplemented with 0.2 mL of chloroform (Sigma Aldrich) and mixed. The samples were then centrifuged (12,000 ×*g*, 15 min, 4 °C), and the aqueous layer was transferred to new tubes. RNA was precipitated by combining 500 μL of isopropanol with the aqueous layer, followed by incubation at room temperature for 10 min and centrifugation as described above. Total RNA was extracted from the supernatant using the PureLink^®^ RNA Mini Kit (Thermo Fisher Scientific, Waltham, USA) according to the manufacturer’s instructions. The extracted RNA was used for quantitative real-time PCR (qPCR). Reverse transcription and reverse transcription quantitative PCR (RT-qPCR) were performed using ReverTra Ace qPCR RT Master Mix with gDNA Remover (Toyobo, Osaka, Japan), THUNDERBIRD^TM^ SYBR qPCR MIX (Toyobo), and Applied Biosystems^®^ 7500Fast (Thermo Fisher Scientific). We assessed the colonization of *H. cinaedi* by measuring the expression of *cdt* at the transcriptional level. Primers for PCR and qPCR are listed in Table S2.

### Histology and fluorescence images

Colons from mice were harvested and fixed with a 4% paraformaldehyde phosphate buffer solution (FUJIFILM Wako). Sections obtained from fixed specimens were permeabilized and incubated with an anti-*H. cinaedi* antibody (1:1,000) as the first antibody at room temperature for 1 h. The anti-*H. cinaedi* antibody was obtained by immunizing rabbits with heat-inactivated *H. cinaedi* MRY08-1234 as an antigen, followed by protein A-purification. The sections were then rinsed and stained with Alexa Fluor 488 (*H. cinaedi*) and 4’,6-diamidino-2-phenylindole (nuclei). Fluorescence signals were visualized using an automated slide scanning system (Olympus VS200, Olympus, Tokyo, Japan) and images were analyzed using ImageJ ver.1.53t.

### Statistical analysis

Differences in the recovery of *H. cinaedi* by culture from infected mice were analyzed using Fisher’s exact tests. For other experiments, the Mann–Whitney test was used for comparison. All statistical analyses were performed using GraphPad Prism version 9.4.1 and differences were considered statistically significant when the *P*-value was <0.05.

## ACKNOWLEDGMENTS

This work was supported by JSPS KAKENHI Grant Numbers 16K18928, 19K07074 (E. R.) and 22K15465 (S. A.), the GSK Japan Research Grant 2017 (E. R.) and by AMED JP20fk0108148 (E. R.).

