## Supplementary material for "Characterization of HcaA, a novel autotransporter protein in *Helicobacter cinaedi*, and its role in host cell adhesion": Table S1 to S2, Fig. S1 to S8

**Table S1. Oligonucleotide primers used for construction of deletion mutants and expression vectors**

| Primer | Oligonucleotide (5′ to 3′)* |
|---|---|
| Generation of deletion mutants of *hcaA* in *H. cinaedi* | |
| Upstream-F | CTTGTAGTTGCTGCCTGCAC |
| Upstream-R | ACTATATCATAAATCTATCCACTTTTCAATCTATATCACGGCTCACACAAGCTAACAGACG |
| aph-F | CGTGATATAGATTGAAAAGTGGATAGA |
| aph-R | GCAGGACGCACTACTCTCG |
| Downstream-F | GCGCACTTCTATACTCTCTGTCGAGAGTAGTGCGTCCTGCAGCGAATCTACGCAAACACA |
| Downstream-R | TACCACAACGCATTCCACTC |
| Generation of expression vectors of HcaA and mutant of the RGD motif of HcaA | |
| RAD-F | TCAAGAG**C**CGATATGCCACAAGAATCT |
| RAD-R | CATATCG**G**CTCTTGACTCAAACTTCAA |

*, Mutated nucleotide is written in bold.

**Table S2. Oligonucleotide primers used for confirmation of knockout of *hcaA* and detection of *H. cinaedi***

| Primer | Oligonucleotide (5′ to 3′) | Target gene (Accession No.) |
|---|---|---|
| Confirmation of knockout of *hcaA* | | |
| KO_check-F | CGCCAAGACTTCGCACGACTTC | *hcaA* (AP017374) |
| KO_check-R | ATCCCACTATCTGCGCCGATTGTG | |
| Detection of *H. cinaedi* in animal experiment by PCR | | |
| cdt-F | GATTTTAATCGTAGCCCTGCG | *cdt* (AP017374) |
| cdt-R | CGCACTCAAACATTCATTGG | |
| Detection of *H. cinaedi* in animal experiment by qPCR | | |
| cdtB_qPCR-F | TACACCATATCCGGACGAGAGC | *cdt* (AP017374) |
| cdtB_qPCR-R | TGGGTCTTTGCCACAAACGG | |

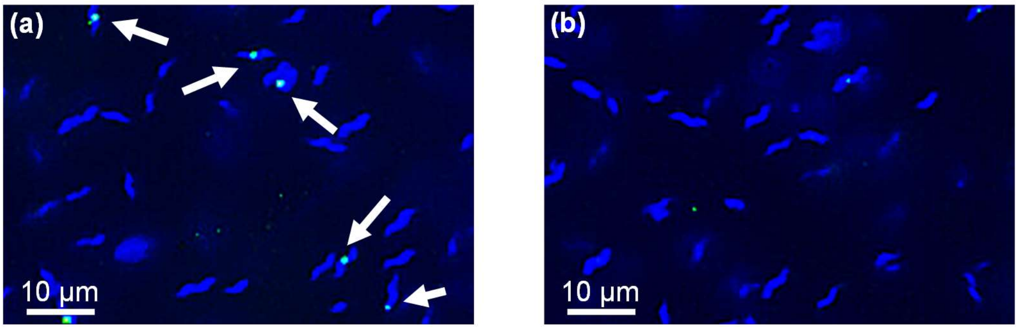


**Fig. S1. HcaA is present on the surface of *H. cinaedi*.**
Image of (a) wild-type MRY08-1234 and (b) HcaA-knockout MRY08-1234. Bacteria were collected and fixed with Blocking One Histo (Nakarai, Kyoto, Japan). The pellets were incubated with an anti-*H. cinaedi* antibody (1:50) at room temperature for 1 h. The pellets were then rinsed and stained with Alexa Fluor 488 (green, *H. cinaedi*) and 4',6-diamidino-2-phenylindole (blue, nuclei). The fluorescence signals were visualized using an inverted fluorescence microscope (Olympus IX83, Olympus, Tokyo, Japan). Fluorescence staining showed the presence of HcaA on the surface of *H. cinaedi* (arrows)

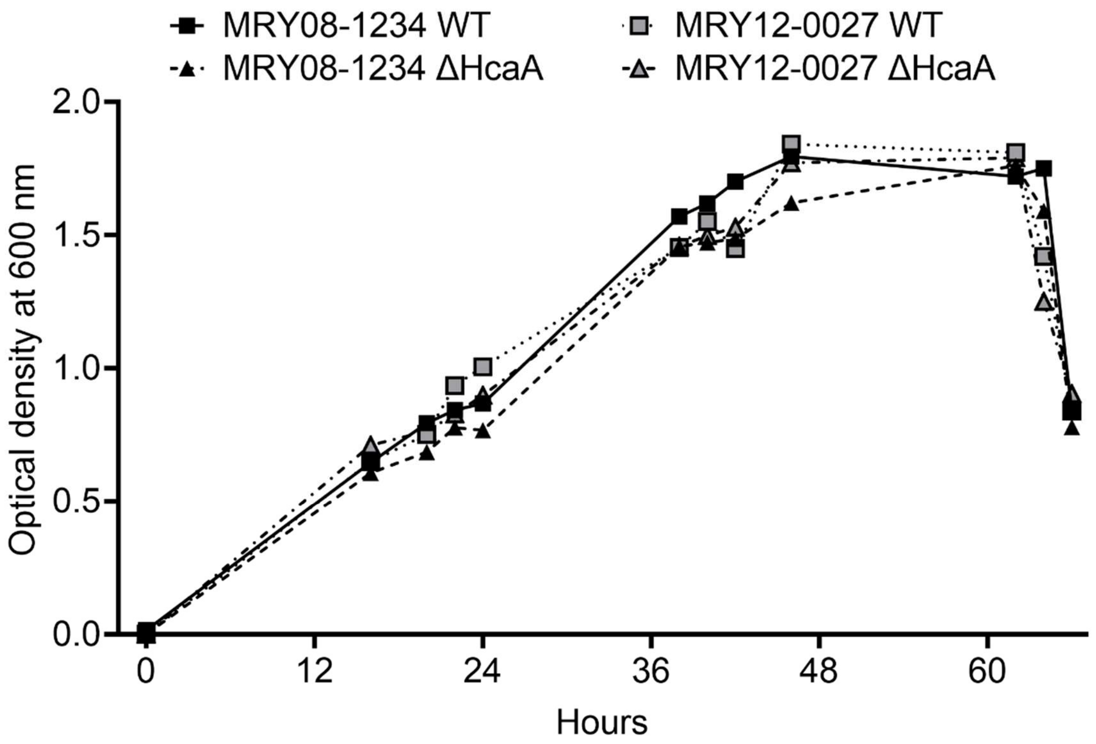


**Fig. S2. Influence of *hcaA* knockout on the growth of *H. cinaedi* MRY012-0027 and MRY08-1234**

The growth potential of wild-type strains and HcaA-knockout strains was measured. No growth defects by knockout of *hcaA* were observed.

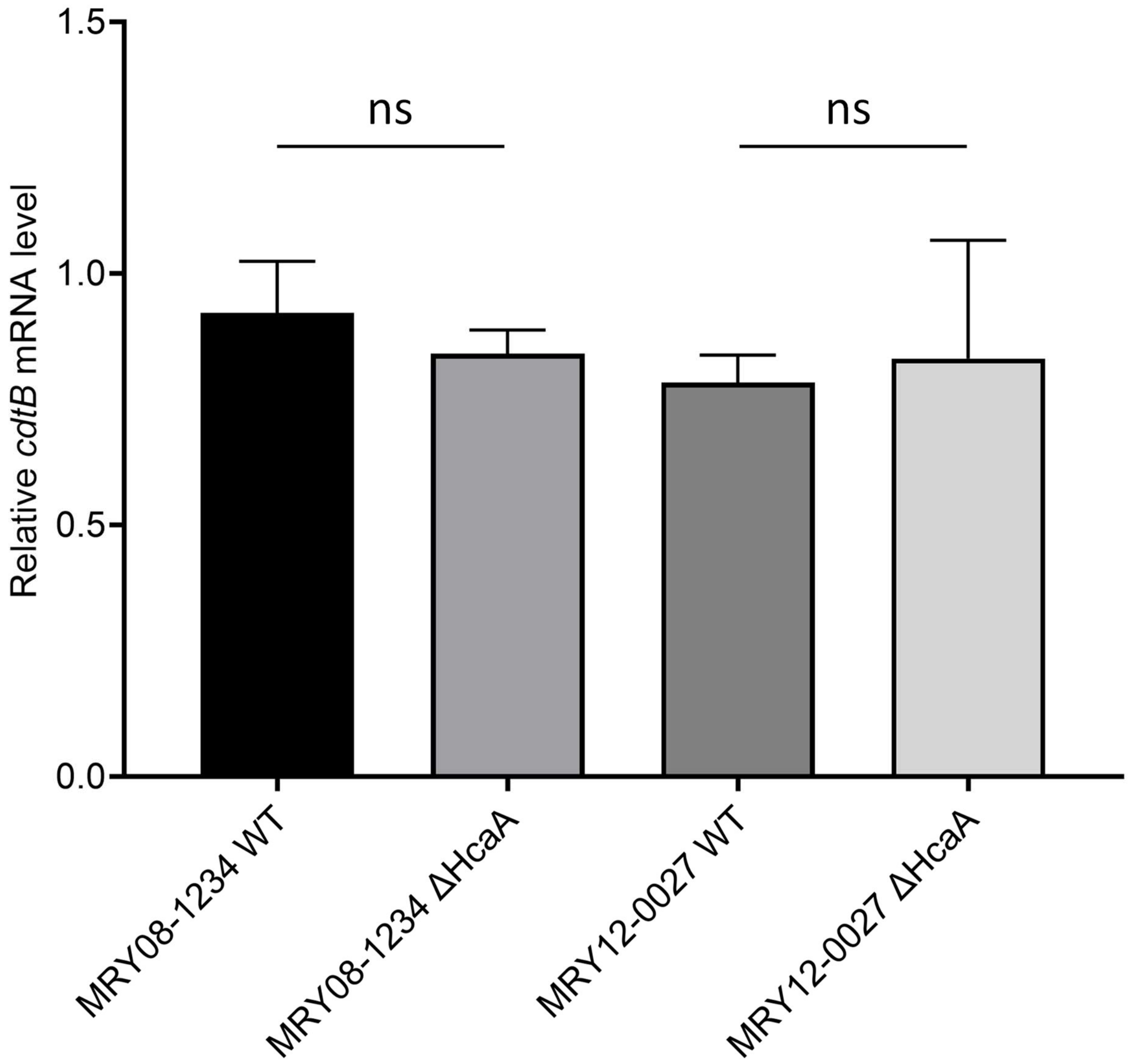


**Fig. S3. No differences in *cdtB* expression levels in wild-type and HcaA-knockout strains**
Relationships between reduced cytotoxicity and *cdtB* expression levels in wild-type and HcaA-deficient strains. Each strain underwent total RNA extraction, reverse transcription, and qPCR following the methods described in the text. The *recA* gene was used as an internal standard (recA_qPCR_F; 5′-CGCACCGCCATTTTAGAGAG, recA_qPCR_R; 5′-AGCCACFCACCACTTTTATC). The results are shown as the mean and standard deviation of six independent experiments. The graph shows the expression levels of *cdtB* in each strain, with no clear differences observed between the wild-type and knockout strains.

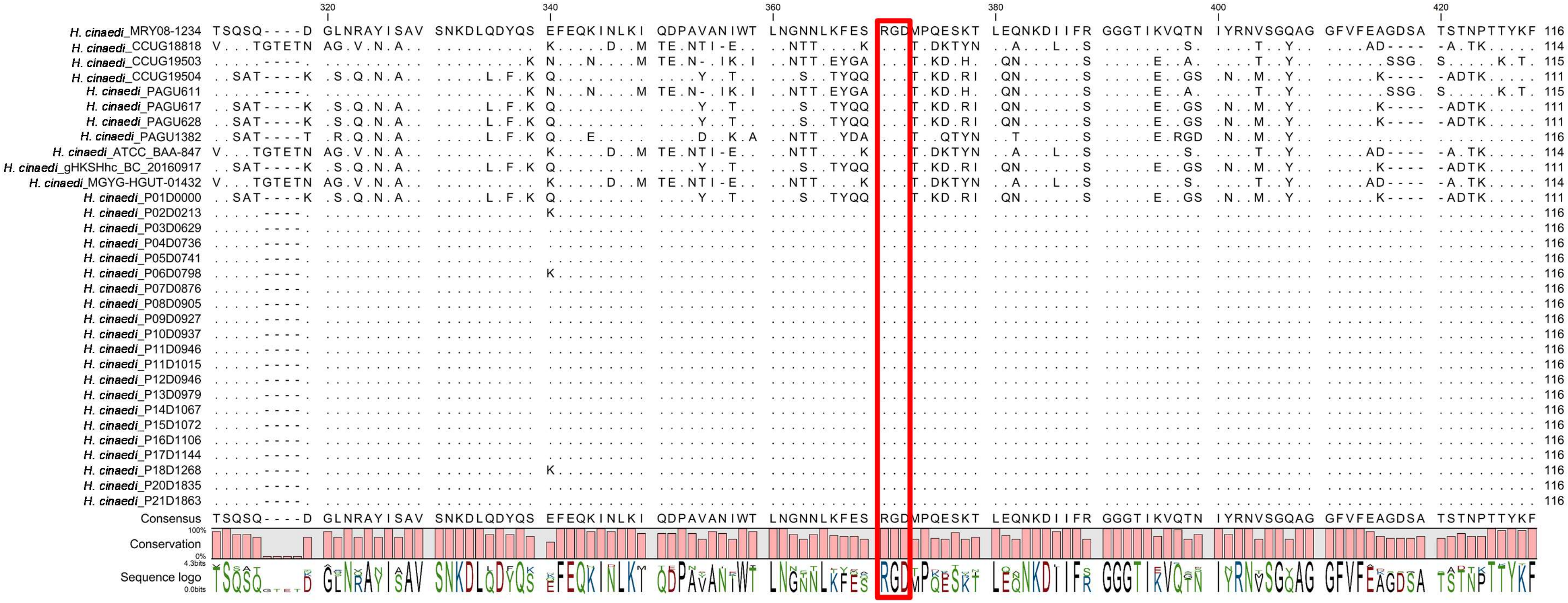


**Fig. S4. Comparison of the partial amino acid sequence of HcaA in various *H. cinaedi* strains.**

All *H. cinaedi* strains possessed HcaA; the RGD motif of HcaA was conserved among strains, as shown in red.

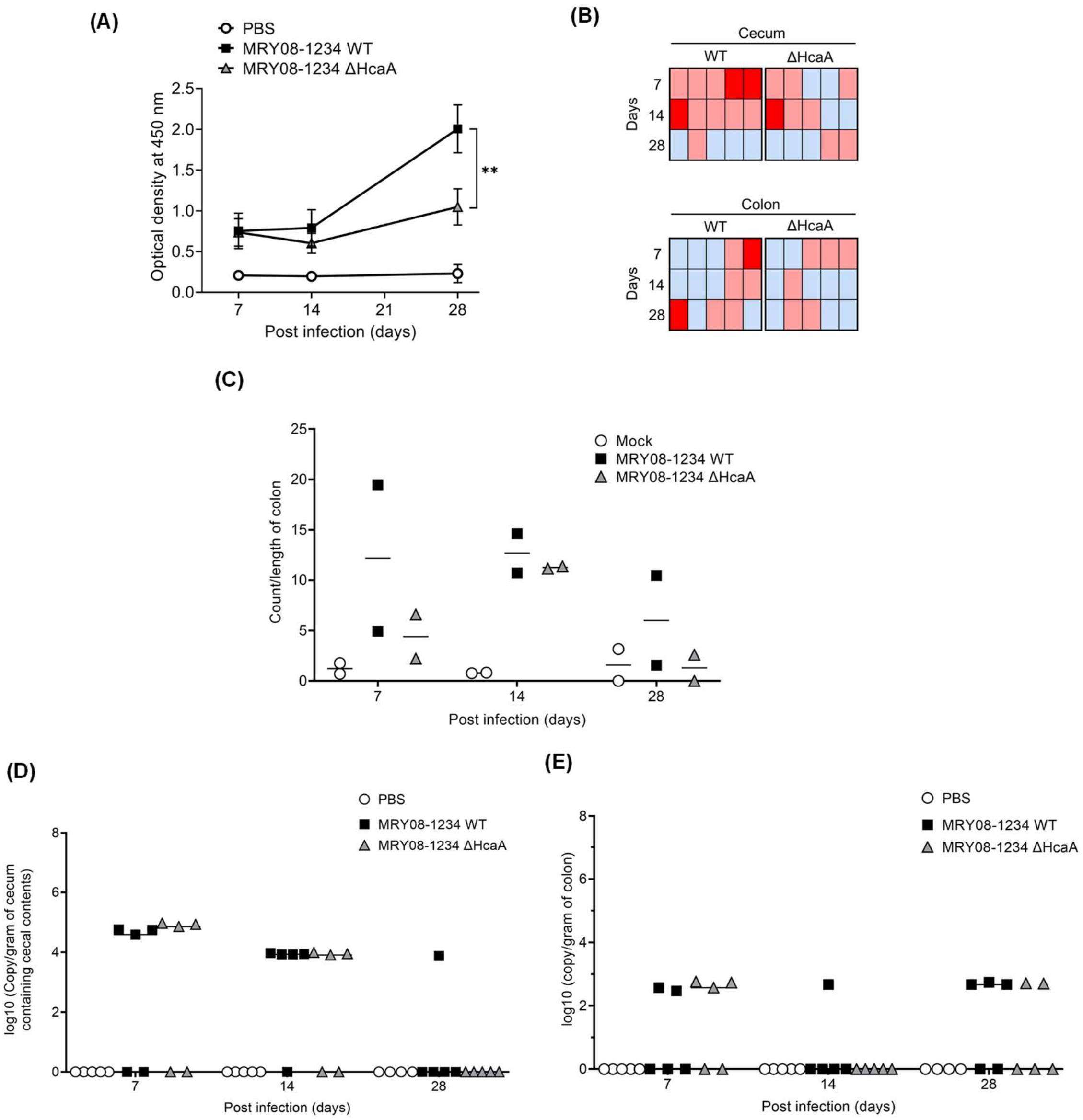


**Fig. S5. Results of the first animal experiment.**

(A) Measurement of anti-*H. cinaedi* antibody titers in infected mice determined by ELISA. Sera from mice at 7, 14, and 28 days after inoculation were used. Squares and triangles indicate serum from the group of infected wild-type (square, MRY08-1234 WT) or HcaA-knockout (triangle, MRY08-1234 ΔHcaA) strains, respectively. Means and SD reflect values for five mice per group. **, $P < 0.01$. (B) Heatmap combining the results of the culture method and qPCR assays. Columns indicate each mouse and rows indicate the duration of infection. Positive results for both the culture method and qPCR are shown in red, positive results for either are shown in pink, and negative results for both are shown in light blue. (C) Quantitative bacteria count per length of colon from *H. cinaedi*-infected mice. Adhesion of *H. cinaedi* to the colon was quantified using the ImageJ ver. 1.53t, by counting the number of *H. cinaedi* in images taken using the 20× objective lens. The results are expressed as numbers of *H. cinaedi* per length of colon. Means and SD reflect values for two mice per group. (D) and (E) To quantify bacterial colonization, correlation curves were generated between Ct values and gene copy numbers. This correlation was utilized to convert Ct values into log10 copies/gram of tissue. The means and standard deviations represent values from five mice per group. Results for the cecum containing cecal contents are indicated by (D), while (E) represent results for the colon.

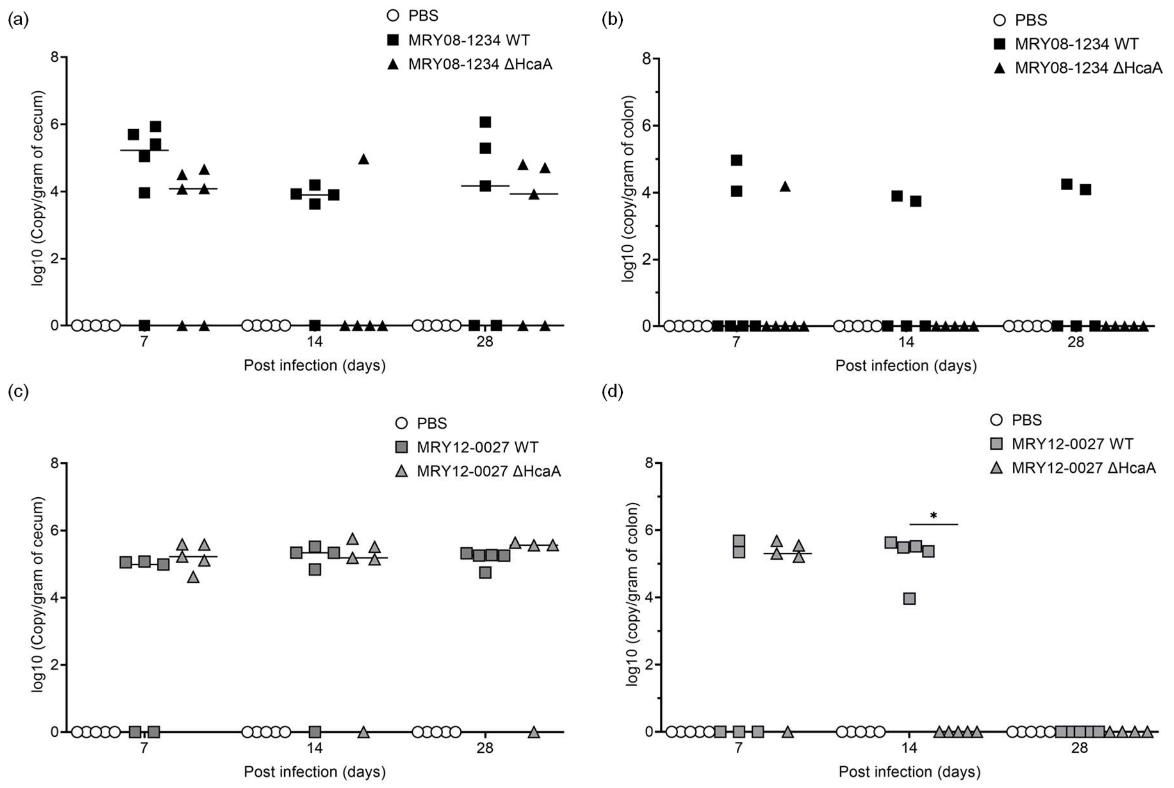


**Fig. S6. Bacterial gene copies per gram of tissues for 7, 14, and 28 days after infection.** To quantify bacterial colonization, correlation curves were generated between Ct values and gene copy numbers. This correlation was utilized to convert Ct values into log10 copies/gram of tissue. The means and standard deviations represent values from five or six mice per group. Results for the cecum are indicated by (a) and (c), while (b) and (d) represent results for the colon. The upper panels show the results for MRY08-1234, and the lower panels show the results for MRY12-0027. *, $P < 0.05$.

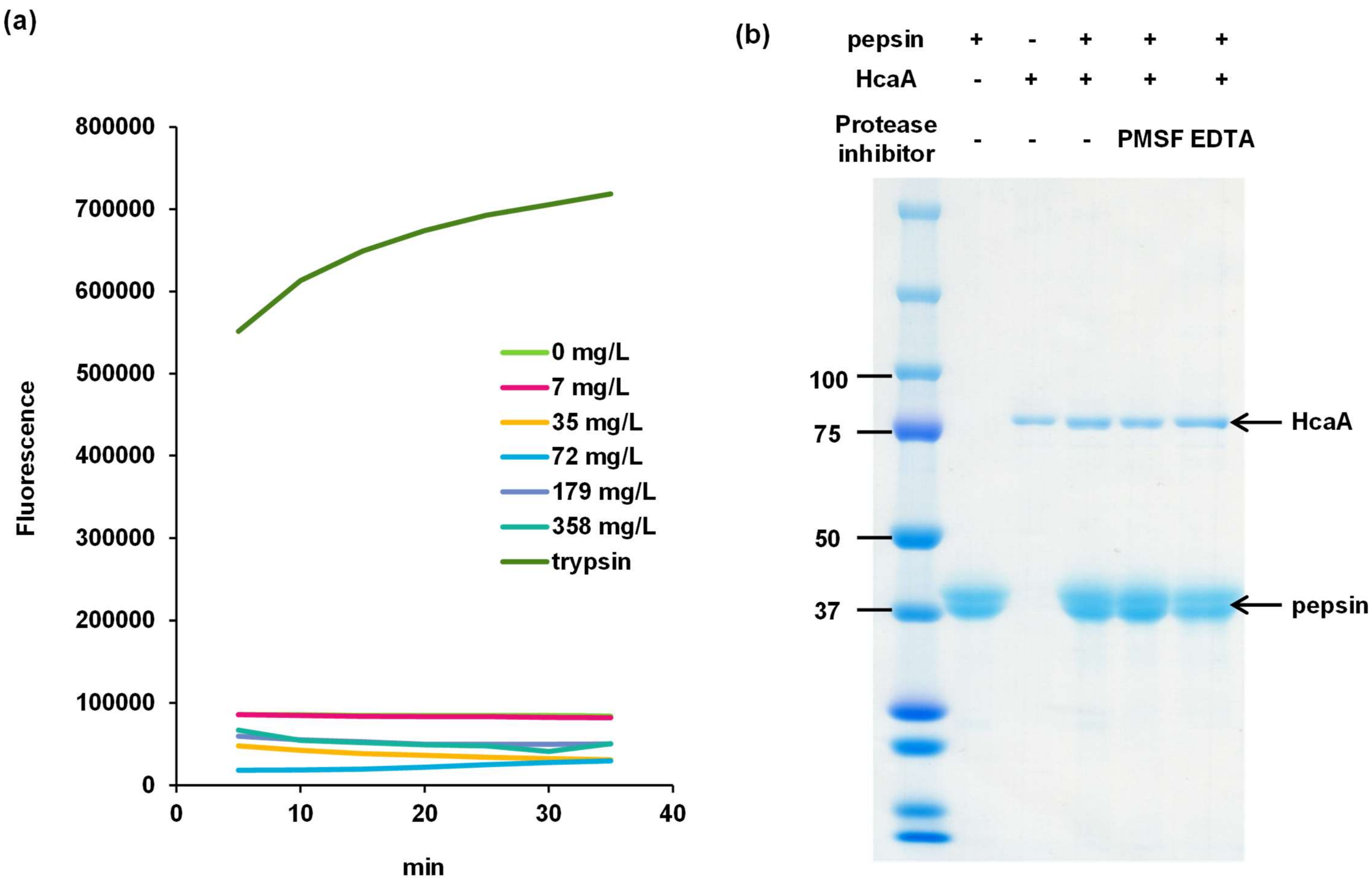


**Fig. S7 Protease activity assays for HcaA.**

(a) Protease activity against casein. Activity was measured using an Amplite$^{TM}$ Universal Fluorimetric Protease Activity Assay Kit, Green Fluorescence (AAT Bioquest, Sunnyvale, CA, USA). HcaA corresponding to positions 31 to 774 (80 kDa) of *H. cinaedi* MRY08-1234, which covers residues that are highly conserved in the superfamily of peptidase S6 protein (e.g., EspP produced by enterohemorrhagic *E. coli*), was cloned into pCold I and purified by HisTrap followed by size exclusion chromatography, as described in the Materials and Methods. The protease activity of HcaA and trypsin as positive controls against fluorescent casein conjugates was measured using the fluorescence signal. The samples were analyzed in duplicate at HcaA concentrations between 0 and 358 mg/L. No protease activity of HcaA was observed in casein. (b) Protease activity of HcaA against pepsin. The purified HcaA was incubated with pepsin at 37 °C for 1 h. The degradation of pepsin was evaluated by SDS-PAGE, followed by CBB staining. No pepsin degradation by HcaA was observed.

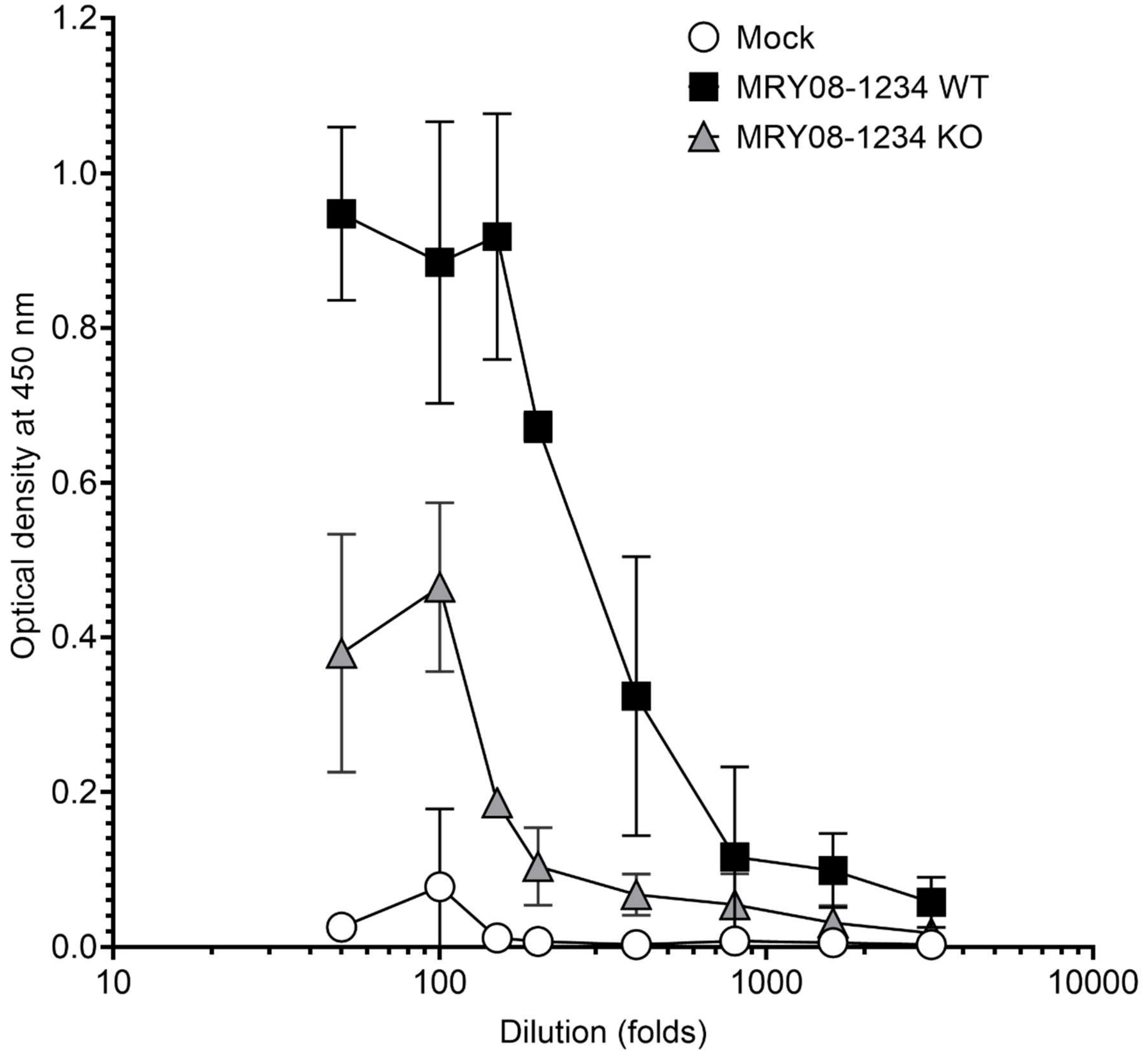


**Fig. S8. Measurement of anti-*H. cinaedi* antibody titers with serial dilution.**
To determine the dilution of the serum, the serum samples from the first animal experiment were utilized for evaluation. The sera were prepared in 2-fold dilutions (50- to 3,200-fold dilutions) and assayed using the same methods described in the manuscript. Sera obtained at 28 dpi were used for analyses. Based on the results, a 100-fold dilution was determined to be suitable and used for subsequent assays.
